# HEMU: an integrated Andropogoneae comparative genomics database and analysis platform

**DOI:** 10.1101/2023.05.19.541421

**Authors:** Yuzhi Zhu, Zijie Wang, Zanchen Zhou, Yuting Liu, Junpeng Shi

## Abstract

The Andropogoneae tribe encompasses various crops with substantial agronomic value such as maize (Zea mays) and sorghum (Sorghum bicolor). Despite the prevalence in released multi-omics data resources, there is a dearth of comprehensive, tribe-level integration and multi-layer omics dataset platform within the tribe, assisting inter- and intra-species comparative analysis from a multi-omics aspect. Here, we first collected a comprehensive atlas of multi-omics datasets within the tribe, including 75 genomes from 20 unique species, transcriptomes from 4,747 samples comprising more than 50 tissues, epigenome data from 90 ChIP-seq samples and 39 ATAC-seq samples, as well as transposable element (TE) annotation for all the genomes. Then, an integrated database and analysis platform, HEMU (http://shijunpenglab.com/HEMUdb/), was constructed. HEMU comprises six sophisticated toolkits, namely genome analysis toolkit, transcriptome-derived analysis toolkit, gene family analysis toolkit, transposable element (TE) analysis toolkit, epigenome analysis toolkit and miscellaneous analysis toolkit, facilitating convenient inter- and intra-species comparative analysis taking advantage of the multi-omics data. Three case studies substantiated the capability of HEMU in conducting gene-centered analysis, transcriptome derived analysis and gene family analysis from a both multi-omics and comparative perspective. In a nutshell, HEMU lowers the barrier of traditional code-based analysis workflow, providing novel insights into modern genetic breeding in the tribe Andropogoneae.

## Introduction

Tribe Andropogoneae within the family Poaceae includes several species of agronomic value, including the C4 crop sorghum (*S. bicolor L. Moench*), maize (*Z. mays L.*) and sugarcane (*S. officinarum L.*), which make up the majority of the world crops production. Despite its significance in agriculture, Andropogoneae also comprises grasses like *Miscanthus sinensis* and *Miscanthus sacchariflorus* that are being investigated as potential biomass crops for renewable energy production (Chupakhin et al., 2022).

With the understanding of the genetic elements and the advancement of sequencing techniques, researchers have been getting insight of the evolution of the genomes, the dynamic of the transcriptomes and the variation of the epigenomes in Andropogoneae. Comparative genomics has shown that the Andropogoneae shared similar physiology while being tremendously genetically diverse, harboring a broad range of ploidy levels, structural variation, and transposons (Song et al., 2021). As a predominant structural element of Andropogoneae genomes, Researchers studied TE (transposable element) insertions across the whole genome in a maize diversity panel and discovered that TE insertional polymorphisms were tagged by SNP markers associating with agricultural trait (Qiu et al., 2021). What’s more, it is also investigated that TE-induced phenotypic changes were associated with domestication and/or diversification in Andropogoneae (Ramachandran et al., 2020).

To understand the molecular mechanism and regulatory network underlying the agricultural and biomass features among tissues and species, researchers also investigate the transcriptome and epigenetic of crops, besides the association and diversity of genome structure. As a result, more and more high-quality genomes for Andropogoneae species have been sequenced and assembled, along with an equivalent amount of RNA-seq, ChIP-seq and ATAC-seq data. The integration and association of omics data, however, cannot be adequately and comprehensively explained from individual study (Suwabe and Yano, 2008). In order to integrate the bioinformatic resources, numerous databases have been built based on species and sequencing types.

Currently, genomic data for species in the tribe Andropogoneae has been compiled in numerous databases, and several bioinformatics tools have been made available to extract new biological information beyond individual plant datasets. With the genomes of several maize inbred lines sequenced, MaizeGDB offers online study of multiple maize genomes, RNA-seq, proteomics, and synteny data. (Portwood et al., 2018) Its omics data, however, is relatively insufficient, with only 6 datasets related to the qTeller mapping on the B73v4 reference genome for gene expression data comparison and analysis. MOROKOSHI uses FL-cDNA and public RNA-seq data to generate expression profiles for each sorghum gene and can visualize gene co-expression networks for users, but only three tissue specific analyses are available, and it is insufficient in expression visualization tools and gene functional annotation functions. (Makita et al., 2014) PlantRegMeg, a plant genetic element database, has integrated transcription factor and *cis*-element information for 63 representative plants and developed an algorithm to screen for functional regulatory elements and interaction. (Tian et al., 2019) Nevertheless, PlantRegMeg is inconvenient for investigating specific regulatory mechanisms associating with transcription, chromatin accessibility, and methylation since it lacks multi-omics data. These databases offer data sources for multi-omics or genetic elements in specific field respectively, but there is a dearth of comprehensive integration and multi-layer omics dataset platform for Andropogoneae, to enable more systematic analysis, sophisticated understanding of the interested genes and genetic element without switching to different databases. Here, we present HEMU (http://shijunpenglab.com/HEMUdb), the first integrated yet handy

Andropogoneae comparative genomics database and analysis platform encompassing six sophisticated toolkits. HEMU enables researchers to easily utilize multi-omics data and perform customized comparative analysis from novel aspects among Andropogoneae species.

## Results

### Overview of multi-omics datasets in HEMU

HEMU encompasses an extensive multi-omics dataset **(Figure 1A, Supplementary Figure S1)**. Firstly, a total of 75 published genomes comprising 20 unique species, including both well-studied model species and recently published non-model species, were utilized with the aim of establishing a comprehensive genomic atlas within the Andropogoneae tribe **(Supplementary Table S1)**. Specially, due to lack of genomic annotation or unavailability of data in certain genomes, a standardized workflow was constructed, assisting the annotation of genomes from certain non-model species. As a result, 5 genomes from varied non-model species were annotated upon the initial HEMU release **(Supplementary Table S2)**, and the number will continue to grow.

**Figure 1.**
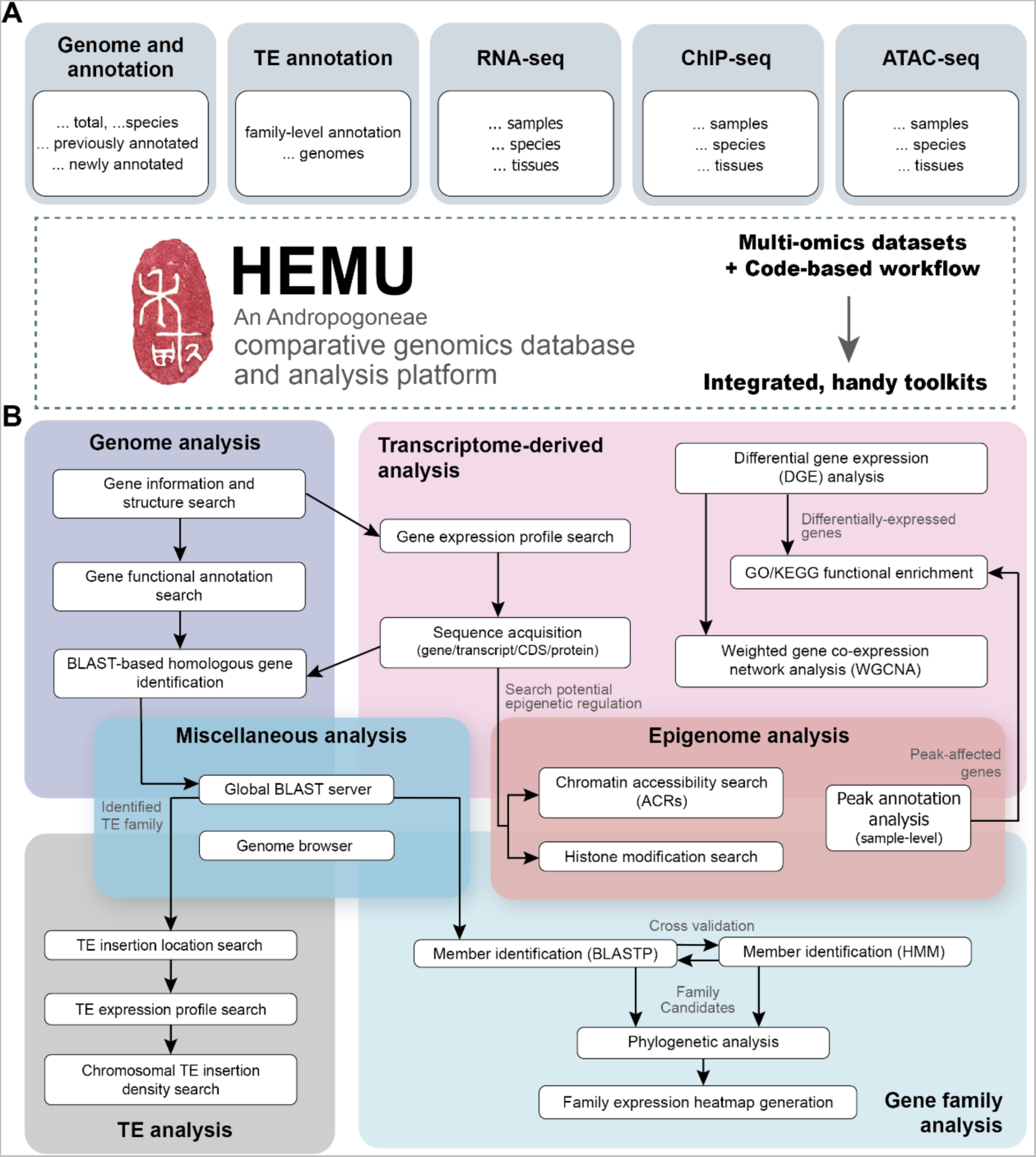
Overview regarding the structure of HEMU database and analysis platform. **(A)** Genomes, transposable element (TE) annotation and multi-omics datasets used to construct HEMU. **(B)** Six main analysis toolkits of HEMU and corresponding modules inside. Arrows indicate potential workflows than can be used to conduct intra- and inter-toolkit analysis.

For transcriptomic data, a total of 4,718 RNA-seq samples from published datasets in the tribe Andropogoneae were meticulously curated. In particular, 1,527 for *Zea mays*, 1,428 for *Sorghum bicolor*, 1,226 for *Saccharum spontaneum*, 338 for *Miscanthus lutarioriparius*, 117 for *Coix lacryma-jobi*, 54 for *Miscanthus sinensis*, 13 for *Microstegium vimineum*, 12 for *Themeda Triandra*, 2 for *Cymbopogon flexuosus* and 1 for *Hyparrhenia diplandra*. Details regarding sample tissue, treatment, accession ID and other auxiliary information can be found in **Supplementary Table S3**. For epigenomic data, a total of 90 ChIP-seq and 37 ATAC-seq samples from published and in-house datasets in the tribe Andropogoneae were collected and processed **(Supplementary Table S4)**. The 90 high-quality ChIP-seq datasets comprise 15 tissues and 11 representative antibodies for histone modification. The 39 ATAC-seq datasets contain 14 unique tissues, laying an excellent foundation for investigating chromatin accessibility.

Notably, transposable element (TE) annotation of all the 75 genomes were also integrated to our multi-omics portfolio, assisting cross-dataset analysis. Moreover, TE expression profiles were estimated based on previously-described RNA-seq datasets, providing a novel perspective to exploring TE-mediated transcriptional regulation as well as potential TE-gene interactions.

### Overview of analysis toolkits in HEMU

With the aim of taking the most advantage of our multi-omics data and lowering the barrier of traditional code-based analysis workflow, HEMU integrated the conventional analysis pipeline into interactive, easily-accessible toolkits. Upon the initial release, six main toolkits, namely genome analysis toolkit, transcriptome-derived analysis toolkit, gene family analysis toolkit, transposable element (TE) analysis toolkit, epigenome analysis toolkit and miscellaneous analysis toolkit, were constructed, aiding efficient comparative analysis within the tribe Andropogoneae **(Figure 1B).**

#### Genome analysis toolkit

This toolkit was designed to provide users with a glance of structural and functional information regarding genes of interest. The “gene information and structure search” module allows users to search basic information of a gene, including its structural components and coordinates on chromosome. Plots can be generated to visualize structures of different transcripts produced by a gene, aiding users to investigate alternative splicing events. The “gene functional annotation search” module facilitates fast search of gene functional information such as GO and KEGG annotation, enabling researchers to rapidly deduce potential function regarding genes of interest.

#### Transcriptome-derived analysis toolkit

Based on transcriptomic data from Andropogoneae species, this toolkit features a one-step solution for conventional RNA-seq analysis. The “gene expression profile search” module generates interactive plots regarding sample-level and tissue-level gene expression, enabling users to acquire expression profiles of interested genes. The “sequence acquisition” module provides an interface for fetching gene, transcript, CDS and protein sequences with option for searching the canonical sequence only. Next, the “differential gene expression (DGE) analysis” module enables users to perform customized differential analysis on RNA-seq datasets provided by the HEMU platform with the aim of screening differentially expressed genes in certain tissues or treatments. Functional enrichment of differentially-expressed genes (DEGs) can be conducted using the GO/KEGG enrichment module, where interactive bubble plot is generated, aiding users to visualize enrichment status intuitively. Additionally, by means of weighted gene co-expression network analysis (WGCNA) module, co-expression network analysis can be performed to identify co-expressed gene modules and estimate module-trait correlations.

#### Gene family analysis toolkit

This toolkit converts basic procedures in conducting conventional gene family analysis into easily-accessible modules. In the “family member identification” module, both HMM-based and BLASTP-based approaches are provided for rapid identification of interested gene family members in the genome of Andropogoneae species. After member identification, the “phylogenetic analysis” module can be utilized to perform multiple sequence alignment, genetic distance estimation and phylogenetic tree construction using corresponding protein sequences. As for expression pattern examination, the “family expression heatmap generation” module is provided for users to conveniently characterize expression levels of gene family members among different samples.

#### Transposable element (TE) analysis toolkit

This toolkit aims at enabling users to efficiently utilize TE annotation data from Andropogoneae species and connect them to other analyses. As for TE expression, a handy “TE expression profile search” module was developed, in which sample-level and family-level TE expression profiles can be easily acquired in the form of interactive plots. The insertion location search module helps users to search TE insertion status flanking or within certain genes and regions. Moreover, an interactive TE chromosomal insertion density module developed using the shiny framework lets users to visualize superfamily level TE distribution along chromosomes within and among species, facilitating inter-species comparative analysis.

#### Epigenome analysis toolkit

Established upon our curated ChIP-seq and ATAC-seq data, this toolkit was designed for conducting related analysis. The “chromatin accessibility search” module was developed, allowing users to locate accessible chromatin regions (ACRs) flanking or within certain genes and regions throughout various tissues. Similarly, the “histone modification/TFBS search” module enables users to obtain histone modification information in different tissues and predict potential TF-binding sites using pre-published ChIP-seq data. More importantly, users can perform peak annotation process, fetching genome-level ChIP-seq and ATAC-seq peak location enrichment results.

#### Miscellaneous analysis toolkit

In conjunction with the previously mentioned toolkits, HEMU boasts a range of auxiliary modules that enhance its capabilities. Notably, the inclusion of the JBrowse2-based genome viewer module empowers users to visualize genomic annotation, TE annotation, distribution of ChIP-seq and ATAC-seq peaks, etc. Furthermore, a global BLAST server was incorporated, containing gene, transcript, CDS, and protein sequences from species in the Andropogoneae tribe.

### Application of integrated toolkits on HEMU datasets

HEMU presents a series of powerful analysis toolkits that utilizes comprehensive multi-omics datasets. By intersecting different modules, it’s possible to conduct comparative genomics study from a multi-omics perspective in the tribe Andropogoneae within minutes. Here, three case studies were put forward, aiding users to take full use of the toolkits.

***Case Study 1****: Explore structure, position, expression profile, sequence and identify potential orthologues of a Zea mays ARF gene Zm00001d023659* using the HEMU genome analysis toolkit and transcriptome-derived analysis toolkit.

*Zm00001d023659* encodes a *Zea mays* auxin response factor (ARF) which specifically binds to auxin-responsive promoter elements (AuxREs), regulating plant development and growth. Taking advantage of the HEMU genome and transcriptome analysis toolkit, it can be observed that the gene situates at around 17.88Mbp, chromosome 10 in the *Zea mays* B73v4 genome. The gene produces 8 transcripts, among which transcript #7 (Zm00001d023659_T7) is considered as the canonical transcript **(Figure 2B)**. Functional annotation attributed the transcript to GO term GO:0000003, GO:0001101 etc. and KEGG term ko:K14486. When investigating gene expression profiles, it can be found that while the gene expresses in most of the samples (1440/1527, 94%) **(Figure 2F)**, it exhibits tissue-specific expression in ear, embryo and shoot apical meristem (SAM) **(Figure 2D; Figure 2E)**, suggesting active auxin response in these tissues. By means of sequence acquisition module, gene sequence of *Zm00001d023659* can be obtained and subsequently fed into the global BLAST module to characterize potential orthologous genes in other Andropogoneae species **(Figure 2G)**. By conducting nucleotide-nucleotide BLAST (BLASTN), a number of genes, such as *SORBI_3008G096000* in *Sorghum bicolor*, *Sspon.02G0027120-3C* in *Saccharum spontaneum* and *Misin14G087300* in *Miscanthus sinesis* can be identified as potential orthologues, facilitating inter-species comparative analysis downstream **(Figure 2H)**.

**Figure 2.**
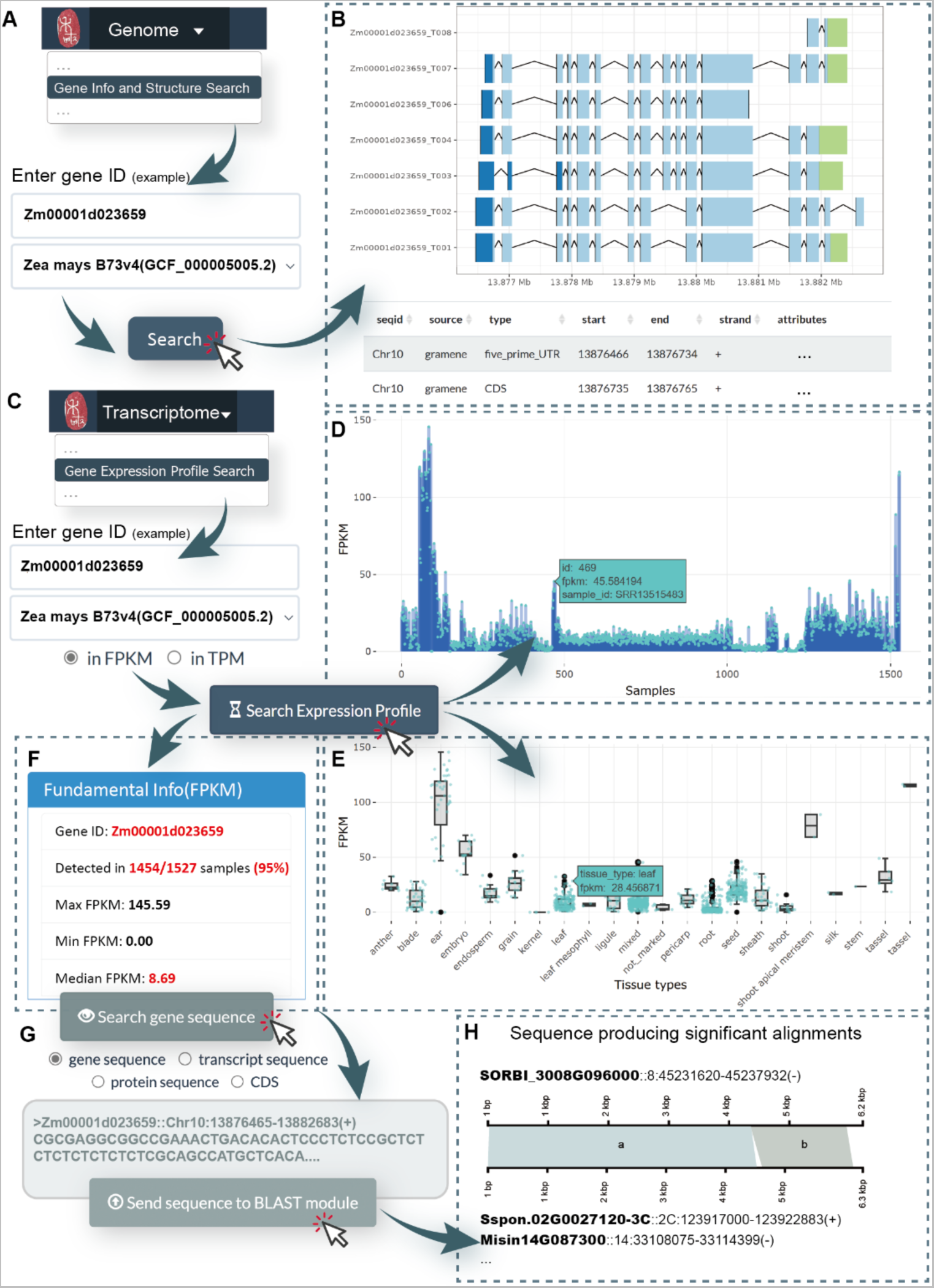
Explore structure, position, expression profile, sequence and identify potential orthologues of a *Zea mays ARF* gene *Zm00001d023659* using HEMU. **(A)** Schematic pipeline of searching gene information and transcript structure in the genome analysis toolkit. **(B)** Basic information and transcript structure of gene *Zm00001d023659*. Deep blue-5’UTR, light blue-CDS, light green-3’UTR. **(C)** Schematic pipeline of searching gene expression profiles in the transcriptome-derived analysis toolkit. **(D-F)** Results generated by searching for *Zm00001d023659* in the “gene expression profile search” module. **(D)** Sample-level expression plot (in FPKM), **(E)** Tissue-level expression plot (in FPKM), **(F)** Panel displaying statistical information regarding expression level among all the samples, in which FPKM>1 was designated as the threshold to identify expressed gene in all samples. **(G)** Schematic pipeline of searching the gene sequence and identify potential orthologs by conducting BLAST against other Andropogoneae genomes. **(H)** Potential orthologs of *Zm00001d023659* identified using BLAST search and graphical overview of aligning regions.

***Case Study 2****: Mining differentially expressed genes in response to heat stress, conduct functional enrichment and constructing co-expressed gene modules in Zea mays* cultivar B73 using the HEMU transcriptome-derived analysis toolkit.

Plant heat stress response (HSR) has been an extensively-studied topic, as excessive temperature negatively affects plant development and metabolism, ultimately impacts yield (Haider et al., 2021; Hatfield and Prueger, 2015). Here, we employed a published RNA-seq dataset (PRJNA396192) of maize in various heat stress conditions (no heat stress, 4h heat stress, 4d heat stress, 4d recovery after heat stress, each with 3 biological replicates) **(Supplementary Table S5)** to demonstrate the ability of HEMU in performing transcriptome-derived analysis including mining differentially expressed genes (DEGs), perform functional enrichment and conducting gene co-expression network analysis. Specifically, a subset derived from *Zea mays* cultivar B73 was used in this case study.

In order to identify genes potentially induced or related to heat stress, differential gene expression analysis was firstly performed with the aid of corresponding module. Two comparisons were designed, namely no heat stress/4h heat stress (0h/4h) and no heat stress/4d heat stress (0h/4d) **(Figure 3A)**. Consequently, a total of 384 and 435 differentially-expressed genes (DEGs) were identified in the 0h/4h comparison and 0h/4d comparison, respectively, based on the default threshold (| log_2_FC |>2 and adjusted P-value<0.05) **(Figure 3B; Figure 3D; Supplementary Figure S2, Supplementary Table S6)**. Principal component analysis (PCA) showed that samples were well clustered in the two comparison groups, indicating alterations in overall gene expression induced by heat **(Figure 3C; Supplementary Figure S2)**.

**Figure 3.**
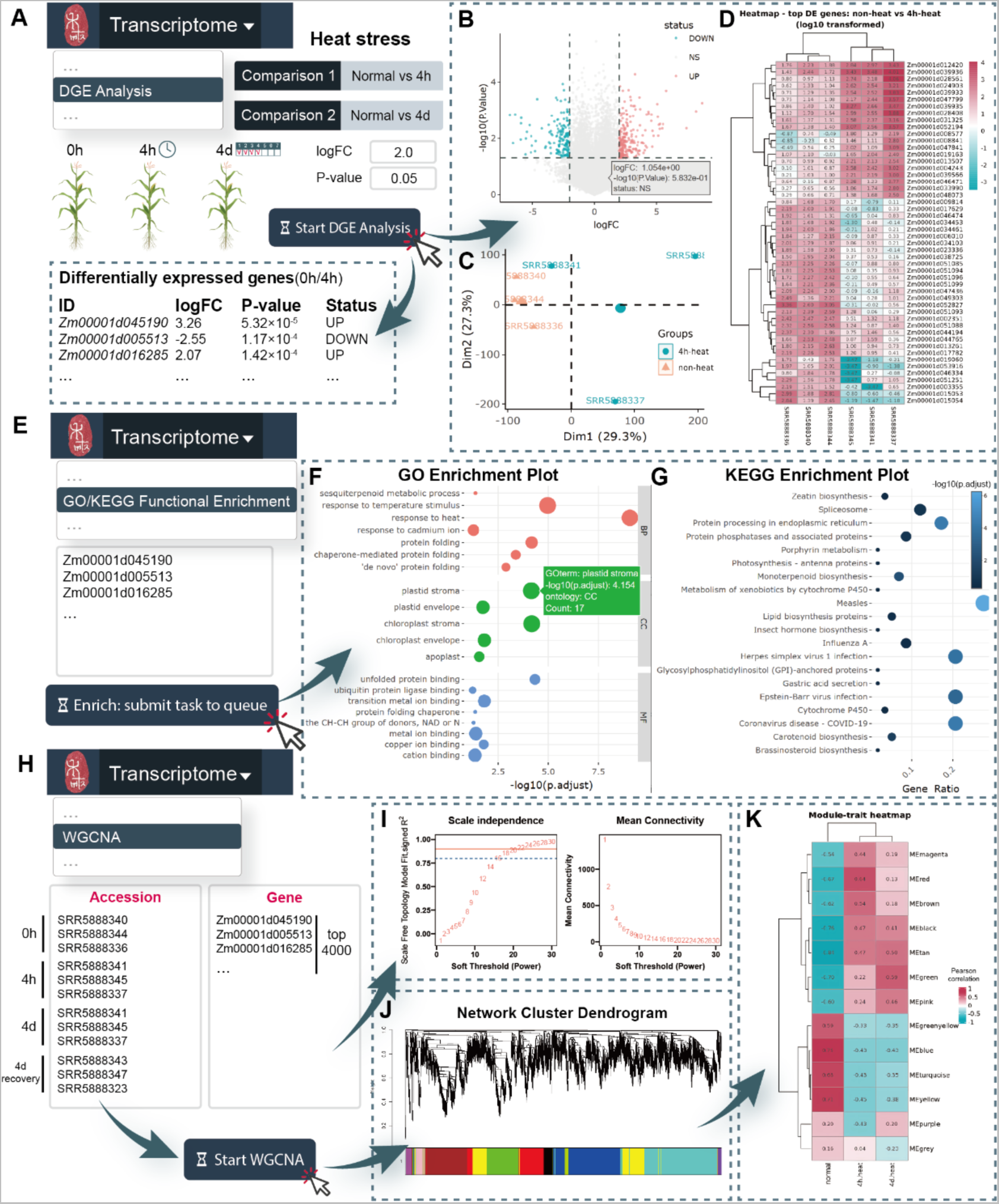
Mining differentially expressed genes in response to heat stress, conduct functional enrichment and constructing co-expressed gene modules in *Zea mays* cultivar B73 using the HEMU transcriptome-derived analysis toolkit. **(A)** Schematic pipeline of conducting differential gene expression (DGE) analysis and an overview of comparisons used in this case study. **(B-D)** Results from the DGE analysis, showing only the 0h/4h comparison. **(B)** Volcano plot of the differentially expressed genes. **(C)** Principal component analysis (PCA) plot of samples from the 0h and 4h heat stress group. **(D)** heatmap regarding log10-transformed TPM values from the top 50 most differentially expressed genes from the 0h/4h comparison. **(E)** Schematic pipeline of performing functional enrichment of differentially expressed genes. **(F-G)** Enrichment results, demonstrated as **(F)** GO enrichment plot and **(G)** KEGG enrichment plot of DEGs in the 0h/4h comparison. **(H)** Schematic pipeline of performing weighted gene co-expression network analysis (WGCNA). **(I-K)** Diagrams generated from the WGCNA module. **(I)** Model scale independence and mean connectivity map with respect to model soft threshold. **(J)** Network cluster dendrogram indicating constructed co-expressed gene modules. **(K)** Heatmap of pearson correlation coefficient between expression levels of co-expression modules and the corresponding treatments.

To determine potential functions of DEGs in *Zea mays* in response to heat stress, GO and KEGG enrichment analysis were conducted on DEGs identified in the two comparisons. In both the 0h/4h and the 0h/4d group, temperature stimulus response and heat response are among the top most enriched GO terms in the Biological Process (BP) section. Notably, in the 0h/4h group, GO terms related to protein folding and binding also exhibit enrichment, suggesting that short-term heat stimulus may result in the formation of misfolded proteins, while part of them then underwent cellular repairing processes **(Figure 3F; Supplementary Figure S3)**. Additionally, plastid (chloroplast) activity was also observed uniquely in the 0h/4h group, which is probably due to the increased energy consumption by protein re-folding. In terms of KEGG pathway enrichment, after removing irrelevant entries such as human diseases (HD), it can be found that spliceosome and endoplasmic reticulum (ER) protein processing pathway demonstrated enrichment in both the 0h/4h and the 0h/4d group, implying that ER-mediated misfolded protein repairing mechanisms and transcription events may constantly take place during the entire heat stress period **(Figure 3G; Supplementary Figure S3)**.

Next, the 4,000 most differentially expressed genes (sorted by | log_2_FC |) were extracted from the two comparisons to construct gene co-expression networks taking advantage of the WGCNA module **(Figure 3H)**. Genes that have FPKM<1 in >30% of the samples were discarded to minimize noise. SFT soft power was chosen based on model recommendations. After co-expression network construction, 13 and 14 co-expressed gene modules were identified in the 0h/4h group and 0h/4d group, respectively **(Supplementary Table S7)**. To discover gene modules that correlate with heat stress, sample-trait correlation analysis was employed. We define modules with a Pearson correlation of r>0.5 between module eigengene (ME) expression level and sample treatment as putative heat-related gene module. As a result, a series of modules correlating with different levels of heat stress were characterized **(Supplementary Figure S4)**. For instance, in the 0h/4h group, the ‘red’ module (ME r=0.64) correlates with short-term heat stress (4h), while the ‘green’ module (ME r=0.59) correlates with long-term heat stress (4d) **(Figure 3K; Supplementary Figure S4)**. Examining eigengenes as well as co-expressed genes within these modules may help elucidating the molecular network underneath. Taken together, these findings are believed to provide reference for further explorations on heat stress response in monocotyledonous plants.

***Case Study 3:*** *Perform comparative analysis on the YABBY gene family in Zea mays, Sorghum bicolor and Coix lacryma, three representative Andropogoneae species, using the HEMU gene family analysis toolkit*.

The *YABBY* gene family encodes a collection of plant-specific transcription factors which were substantiated to participate in the formation of adaxial-abaxial polarity and the regulation of lateral organ development (Ha et al., 2010; Kumaran et al., 2002; Tanaka et al., 2012). The significance of the *YABBY* gene family warrants its utilization as an exemplar for showcasing the competency of HEMU in conducting gene family analysis. In this case, three representative species in the tribe Andropogoneae, *Zea mays* (maize), *Sorghum bicolor* (sorghum) and *Coix lacryma* (job’s tears), were employed to provide a comparative aspect regarding gene family members in near relatives.

In terms of gene family member identification, HMM-based and BLAST-based screening was firstly performed. By means of the gene family identification module, HMM profile of the Pfam *YABBY* domain (PF04690) was firstly used to characterize potential *YABBY* genes in the three genomes (sequence E-value threshold set at 10^-5^, corresponding to a previous study (He et al., 2021)). Candidates were further validated using domain information and protein-protein BLAST with *YABBY* reference sequences **(Figure 4A)**. Consequently, 15 *YABBY* gene candidates were found in *Zea mays*, 8 in *Sorghum bicolor* and 8 in *Coix lacryma-jobi* **(Supplementary Table S8)**.

**Figure 4.**
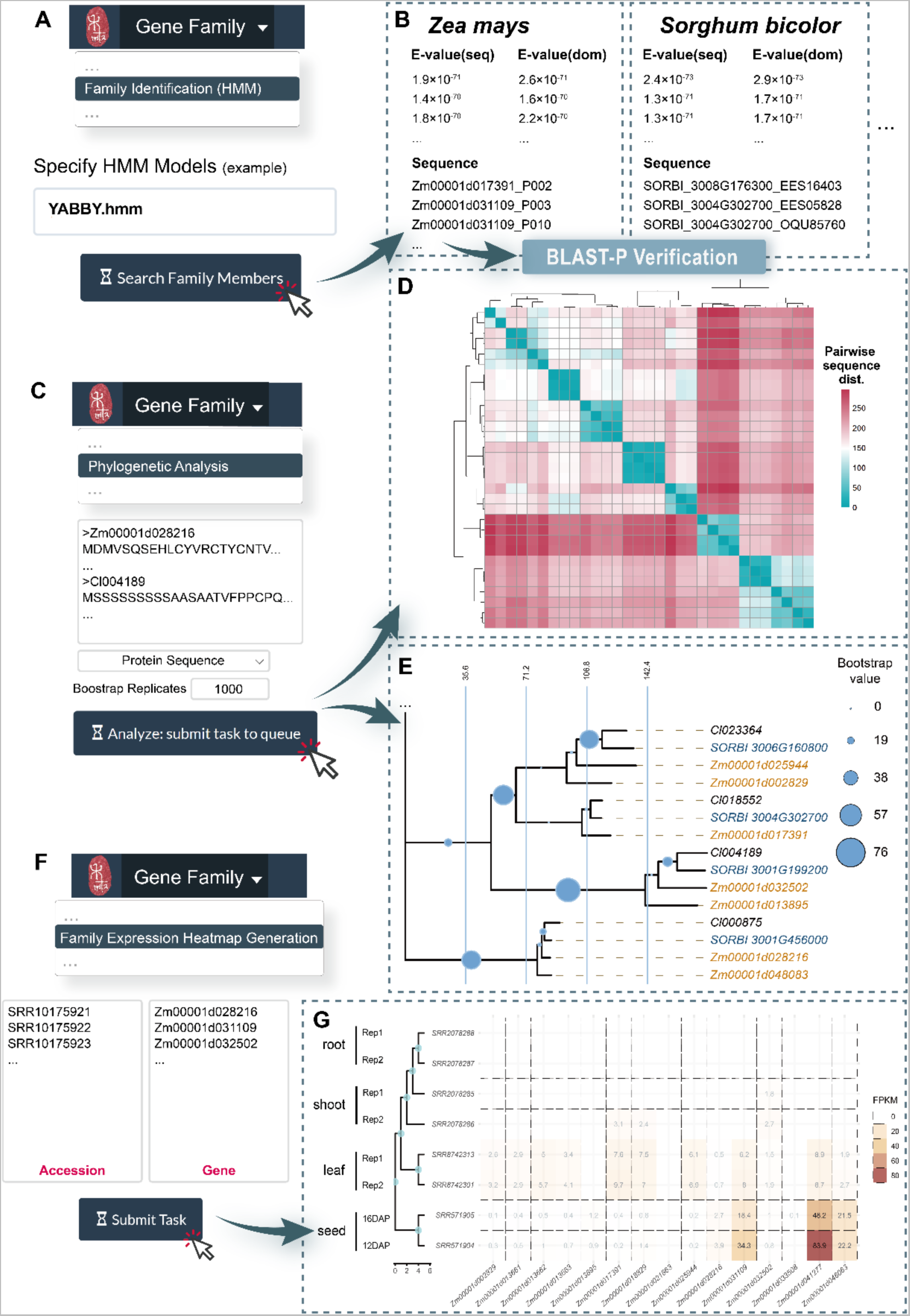
Perform comparative analysis on the *YABBY* gene family in *Zea mays*, *Sorghum bicolor* and *Coix lacryma-jobi*, three representative Andropogoneae species, using the HEMU gene family analysis toolkit. **(A-B)** Identifying putative gene family members in the three species. **(A)** Schematic pipeline to characterize *YABBY* gene family members combining HMM profile and BLASTP. **(B)** Example of identified *YABBY* family members. seq: sequence; dom: domain. **(C-E)** Phylogenetic analysis using protein sequences of identified *YABBY* family members in the three species. **(C)** Schematic pipeline on task submission. **(D)** Pairwise genetic distance (Kimura 2-parameter) heatmap of all protein sequences. **(E)** Part of the phylogenetic tree constructed using neighbor-joining algorithm and testified with 1,000 bootstrap replicates. The full tree is available in supplementary information. **(F)** Schematic pipeline of generating gene family member expression heatmap in the corresponding module. **(G)** Expression heatmap of *YABBY* gene family members in *Zea mays*. Four consistent tissues, namely root, shoot, leaf and seed were chosen to depict the overall expression atlas in the three species. The full heatmap is available in supplementary information.

To further determine hierarchical relationship between these genes, phylogenetic analysis was performed using the corresponding module. Multiple sequence alignment (MSA) of protein sequences demonstrated various conserved regions justified within all the identified candidates **(Supplementary Figure S5)**. Pairwise sequence distance heatmap showed clusters comprising genes from different species, suggesting putative *YABBY* subfamilies among the three species **(Figure 4D; Supplementary Figure S6)**. Next, neighbor-joining dendrogram was constructed, revealing that hierarchical positions of genes were uniformly distributed both within and among species. Taking a closer inspection of the dendrogram, 4-6 potential *YABBY* subfamilies can be potentially characterized combining bootstrap-supported branches (n>50) and the hierarchical information **(Figure 4E, Supplementary Figure S6)**.

Furthermore, expression profiles of *YABBY* family in different tissues were also investigated among the three species using the gene family expression heatmap module **(Figure 3F)**. Comparing heatmap generated from *YABBY* family members generated from the three species, it can be observed that the *YABBY* members generally do not exhibit high expression in most of the tissues, this result somehow agrees with its nature as a tissue-specific transcription factor. Notably, *Zm00001d041277*, *Zm00001d031109*, *Cl035698* and *SORBI_3008G176300*, four *YABBY* genes that form a clade in protein sequence pairwise distance heatmap, display different expression patterns among tissues selected in this case **(Supplementary Figure S7)**. Specifically, *Zm00001d041277, Zm00001d048083* and *Cl035698* show significant expression in seeds, while *SORBI_3008G176300* expresses mostly in roots and shoots, suggesting a potential alteration of *YABBY* gene function following the diversification of *Sorghum bicolor*, despite their relative high sequence similarity.

## Discussion

In this study, we collected and processed Andropogoneae genomes and multi-omics data from various studies and databases, then constructed HEMU, an Andropogoneae comparative genomics database and analysis platform hosting both multi-omics data and sophisticated toolkits. Moreover, three case studies were provided to fully demonstrate the ability of HEMU in assisting genetic breeding. HEMU contains the following features when compared to published datasets pertaining to specific Andropogoneae species.

1. As a multi-omics database, HEMU integrates sophisticated and standardized processing omics data and provides a handy analysis platform. Users can not only query basic information of the genes such as their structures and functions, but can also explore and visualize their differences, dynamics and regulatory relationships through differential expression analysis, GO analysis and epigenomic analysis, etc.
2. As a tribe-scale database, HEMU comprises information of genomic elements for all collected species and provides comparative genomics analysis tools. HEMU originality annotates the most comprehensive TEs of Andropogoneae and provides online analytical tools such as TE expression profile search, insertion location and density search.
3. As an open-source database, all the data analysis and visualization scripts could be obtained directly from HEMU. The codes of the entire project, including MySQL database construction, Django framework and User Interface design, are also accessible via GitHub and developers could utilize the framework and build other analysis platforms.

Scientists have proposed upcoming breeding approaches known as 5G breeding (Varshney et al., 2021) and breeding 4.0 (Wallace et al., 2018), which feature the production of large amounts of omics data and breeding information that can be used to find novel genome editing loci and design new breeding strategies efficiently. Andropogoneae, as the tribe with the most agriculturally and energetically valuable crops, includes exceptionally valuable genetic resources for breeding, necessitating large-scale and comparative genomic investigation to meet breeding 4.0 needs. As a result, HEMU is conducted to become an important data center for Andropogoneae crop breeding, despite the fact that the current version still has some limitations, such as insufficient expression and epigenetic data for non-model species and unbalanced sequencing data for various tissues. Further development of HEMU will be focus on automated and interactive data addition workflow, plugin integration, and omics data types expansion. In general, HEMU aims to give researchers and breeders with a comprehensive data sharing platform and useful analytical tools, accelerating and improving Andropogoneae research and breeding jointly.

## Methods

### Identification of transposable elements (TEs)

A tailored annotation workflow was constructed based on previously-reported annotation pipeline (Ou et al., 2019) to accurately characterize TEs in all Andropogoneae genomes catalogued within the scope of this study, Specifically, LTR-retriever (Ou and Jiang, 2018), LTR-finder (Xu and Wang, 2007) and LTRharvest (Ellinghaus et al., 2008) were utilized for discovering class-I LTR retrotransposons, TIR-learner (Su et al., 2019) for the characterization of class-II TIR transposons, and HelitronScanner (Xiong et al., 2014) for finding potential helitrons. Consequently, a structure-based TE library was constructed. Then, RepeatMasker version open-4.1.1 **(see data availability)** was used to map elements from the curated TE library to the original sequence while constructing a genome-scale TE annotation. The 80-80-80 rule (Wicker et al., 2007) was then applied to the final library to classify TEs into unique families, wherein the similarity cutoff was set at 80%.

### Annotation of non-model Andropogoneae genomes

Considering variances in data processing and analysis methods across studies may pose difficulty, a unified genome annotation workflow was constructed and implemented upon non-model species in the tribe Andropogoneae that possessed a published genome but lack quality annotation.

For genomes with relatively ambiguous or low-quality annotation, we implemented the MAKER annotation pipeline (v3.01.03). Each genome was annotated using successive rounds of MAKER, combining both *de novo* predictions and homology-based evidences (Cantarel et al., 2008) to form a relatively high quality annotation. Particularly, repeat regions within the genome were first masked referring to the previously constructed TE library. Then, biological data were provided to EvidenceModeler (Haas et al., 2008) for the initial run. With respect to protein evidences, we selected the UniProt protein database (Bateman et al., 2021) as well as proteins from the well-annotated *Zea mays* cultivar B73 (Jiao et al., 2017), which shares substantial homology to non-model Andropogoneae species in this study. Transcript evidences was provided as transcriptomes assembled by Trinity (v2.1.1) (Haas et al., 2013) using published RNA-seq data corresponding to the certain species. Augustus (Stanke et al., 2006) and SNAP (Korf, 2004) were used as *ab initio* predictors for potential gene models. Benchmarking Universal Single-Copy Orthologs (BUSCO) scores were used to validate integrity and quality of the annotation. All the newly-annotated genomes exhibit complete BUSCOs greater than 85% in embryophyta datasets (n=1614), indicating a relatively good annotation quality.

Gene models with an AED (Annotation Edit Distance) greater than 0.5 were discarded to minimize false positives. Gene ID for genomes of these newly annotated species was given combining the first letter from its generic name, the first three letter from its specific name, and a six-digit unique gene identifier (e.g., *Ttria_000001* for *Themeda Triandra*).

### Transcriptome-derived analysis

FASTP (v.0.20.1) was used primarily in clipping potential contaminant adaptor sequences and filtering out low quality reads (Chen et al., 2018). Read mapping to reference genome was conducted with Hisat2 (v.2.2.1) using default parameters (Kim et al., 2019). Alignment results were subsequently sorted and converted into binary .bam files with samtools (v1.12) (Danecek et al., 2021). Gene expression level was normalized by Stringtie (v2.1.5) (Pertea et al., 2015) in the form of both fragments per kilobase of transcript per million mapped reads (FPKM) and transcripts per kilobase million reads (TPM). Genes with an FPKM or TPM >1 were considered to be expressed in the corresponding samples. Bulk RNA-seq differential gene expression (DGE) analysis was set to be carried out with R package limma (Ritchie et al., 2015) using log_2_(N+1) transformed TPM values. Before transformation, TPM values were first normalized between samples as to minimize bias caused by different library sizes. The default threshold set for identifying differentially expressed genes (DEGs) were | log_2_FC |>2 and adjusted P-value<0.05. Weighted gene co-expression network analysis is chiefly enabled by R package WGCNA (Langfelder and Horvath, 2008). Protein functional annotation was conducted with eggNOG-mapper (version emapper-2.1.9) (Cantalapiedra et al., 2021) based on eggNOG orthology data (Huerta-Cepas et al., 2019). Sequence searches were performed using DIAMOND (Buchfink et al., 2021). GO and KEGG enrichment analysis was configured to be performed by R package clusterProfiler (Wu et al., 2021).

### Gene family identification and analysis

Hidden Markov Models of 19,632 gene families were curated and downloaded from the Pfam database (https://www.ebi.ac.uk/interpro/download/Pfam/) (Mistry et al., 2021). HMMER (v3.3.2) (http://hmmer.org) and BLASTP (v2.13.0+) are integrated to the server backend, assisting the screening of gene family members. For phylogenetic analysis of identified gene families, ClustalW, ClustalOmega and Muscle algorithm were used for multiple sequence alignment, while neighbor-joining based phylogenetic tree construction and bootstrap validation were performed with R package ape (Paradis and Schliep, 2019).

### TE expression level estimation

Taking advantage of the sorted alignment files (.bam) from transcriptome data analysis, TEtranscripts (v2.1.4) (Jin et al., 2015) was subsequently implemented to estimate both gene and TE read counts corresponding to each of the RNA-seq samples. Raw counts for each TE family were then normalized into fragments per kilobase of transcript per million mapped reads (FPKM) and transcripts per kilobase million reads (TPM) based on the combined length of all TE family members, using tailored R scripts **(see data availability)**.

### TE-derived analysis

To visualize chromosomal distribution of TE families, we split each chromosome into customizable 50kbp-1Mbp windows, then calculated TE coverage in each window. Estimation of insertion time for the identified LTR retrotransposons (LTR-RTs) involved computing sequence divergence between the two LTRs located at either end of the element. Using formula T = K/2µ, where K is the divergence between two LTR sequences and µ represents the species-specific substitution rate, it is possible to calculate the insertion date of each LTR-RT, represented in millions of years ago (Mya). The substitution rate used in this study was 1.3 × 10^-8^ substitutions per site per year, as proposed for LTR-RT elements in *Oryza sativa* (Gaut et al., 1996; Ma and Bennetzen, 2004), which share great homology to species in the tribe Andropogoneae.

### Epigenome-derived analysis

In the case of ChIP-seq data processing, identical methods were implemented to clip adaptor sequences and remove low-quality reads as in RNA-seq data analysis. After read filtering, we first map reads back to the genome using bowtie2 (v2.2.8) (Langmead and Salzberg, 2012). Duplicated reads were then removed with Picard MarkDuplicates (v2.27.4) (https://broadinstitute.github.io/picard/). Peak calling was done by macs2 (v2.1.4) (Gaspar, 2018) with the parameters “-f BAM -g 1000000000 -B -p 0.00001 --nomodel --extsize 147 --broad” and the samples within the same antibody and tissue were merged by IDR (v2.0.4.2) (Li et al., 2011). The R ChIPseeker package (Yu et al., 2015) was used primarily for visualizing peak distribution and annotate peak-related genes across the genome.

The same procedures as ChIP-seq data analysis for read filtering and mapping to the reference genome were used in terms of ATAC-seq data. Removal of duplicates was also conducted with Picard MarkDuplicates (v2.27.4) and peak calling by macs2 (v2.1.4). TSS enrichment and visualization was enabled by deeptools computeMatrix (v3.5.1) (Ramirez et al., 2016), while Tn5 filtering was performed using customized bash and R scripts **(see data availability)**.

### Database implementation

HEMU (http://shijunpenglab.com/HEMUdb) is constructed upon on the Django (v3.2) framework with Node.js (v16.18.0) as JavaScript library and Celery (v5.0.5) as handler for backend task asynchronization. Task message broker is enabled by Redis (v7.0.9). The main server instance is hosted over uWSGI (v2.0.20) and runs on a nginx web server (v1.18.0) with MySQL (v8.0.27) as its core database engine. Backend codes dedicated for data curation, statistical analysis and visualization are implemented using Python (v3.7.12) and R (v4.1.3). Peripheral interactive analysis platform is built upon the R Shiny (v1.7.1) framework. The database as well as the analysis platform is available online without requirement for registration.

## Data availability

Online interface of HEMU is publicly accessible at http://shijunpenglab.com/HEMUdb and is open for research use without user registration.

## Code availability

The backend framework of HEMU is released as an open-source project available in the GitHub repository (https://github.com/EdwardZhu02/HEMU-Database). Scripts for generating plots are also available on Github.

## Author contributions

Y.Z. and Z.W. conceived the project and collected publicly available multi-omics data regarding Andropogoneae species. Y.Z., Z.W. and Z.Z. participated in the construction and diagnosis of the analysis platform. Y.L. conducted ATAC-seq data analysis regarding maize samples. Y.Z. and Z.W. wrote the manuscript.

## Acknowledgments

We sincerely thank Zhijian Chen from school of agriculture, Sun Yat-sen University for platform logo designing, and Jun Cai from school of agriculture, Sun Yat-sen University for informative comments and suggestions on the *YABBY* gene family. No conflict of interest is declared by the authors.

## Supplemental information

**Supplementary Table S1.** The comprehensive genomic atlas of tribe Andropogoneae.

**Supplementary Table S2.** Basic information of newly annotated non-model Andropogoneae genomes upon the initial HEMU release.

**Supplementary Table S3.** The comprehensive transcriptome atlas of tribe Andropogoneae.

**Supplementary Table S4.** The comprehensive epigenomic atlas of tribe Andropogoneae.

**Supplementary Table S5.** Detailed sample information and grouping of *Zea mays* RNA-seq dataset PRJNA396192 containing various heat stress conditions.

**Supplementary Table S6.** Information of differentially-expressed genes in *Zea mays* related to heat stress conditions (0h/4h and 0h/4d).

**Supplementary Table S7.** Co-expression gene modules of the maize differentially-expressed genes between induced and related to heat stress conditions.

**Supplementary Table S8.** Information of *YABBY* gene family member candidates in *Zea mays*, *Sorghum bicolor* and *Coix lacryma-jobi*.

**Supplementary Figure S1.**
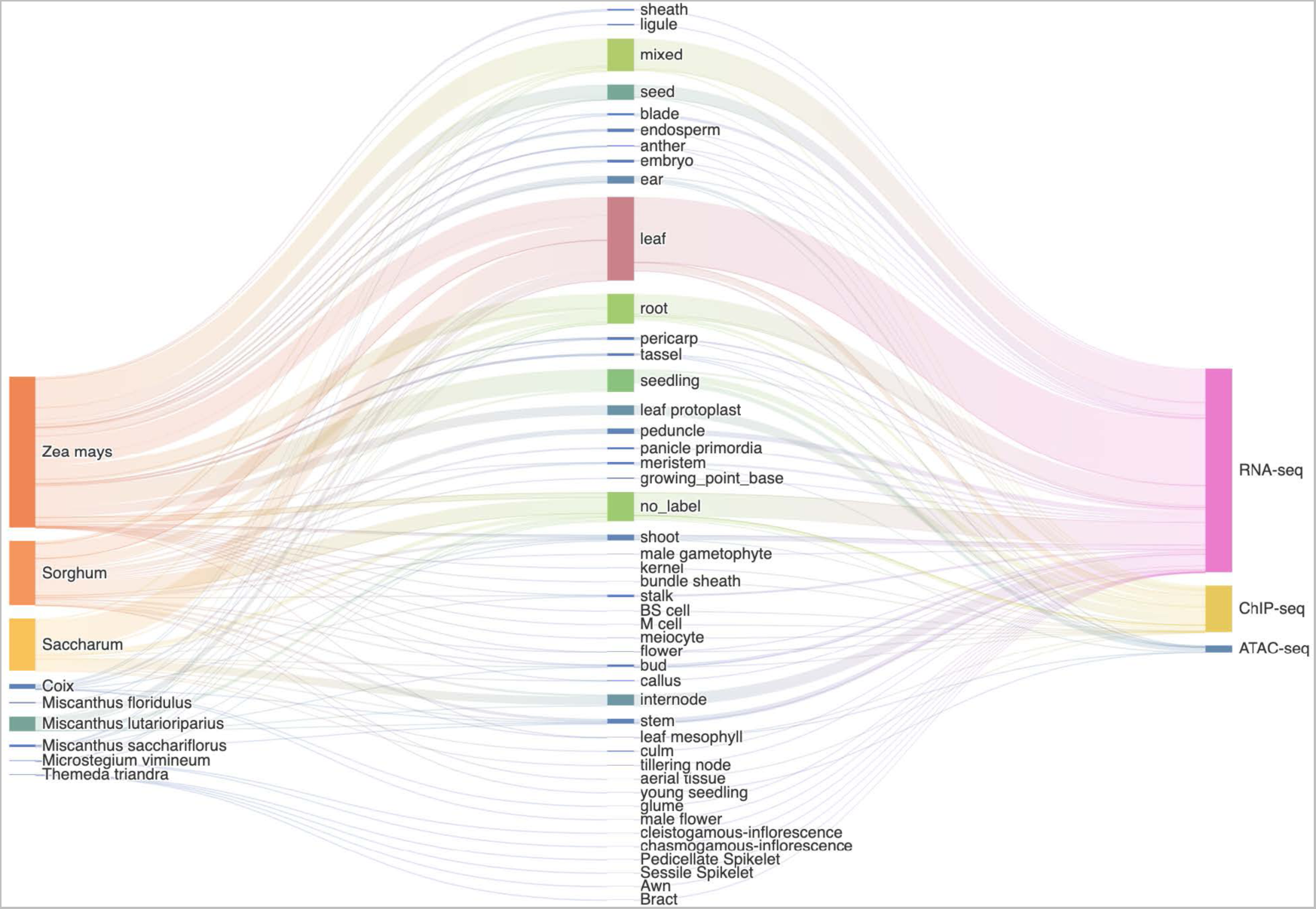
Sankey plot of multi-omics data utilized in the construction of HEMU.

**Supplementary Figure S2.**
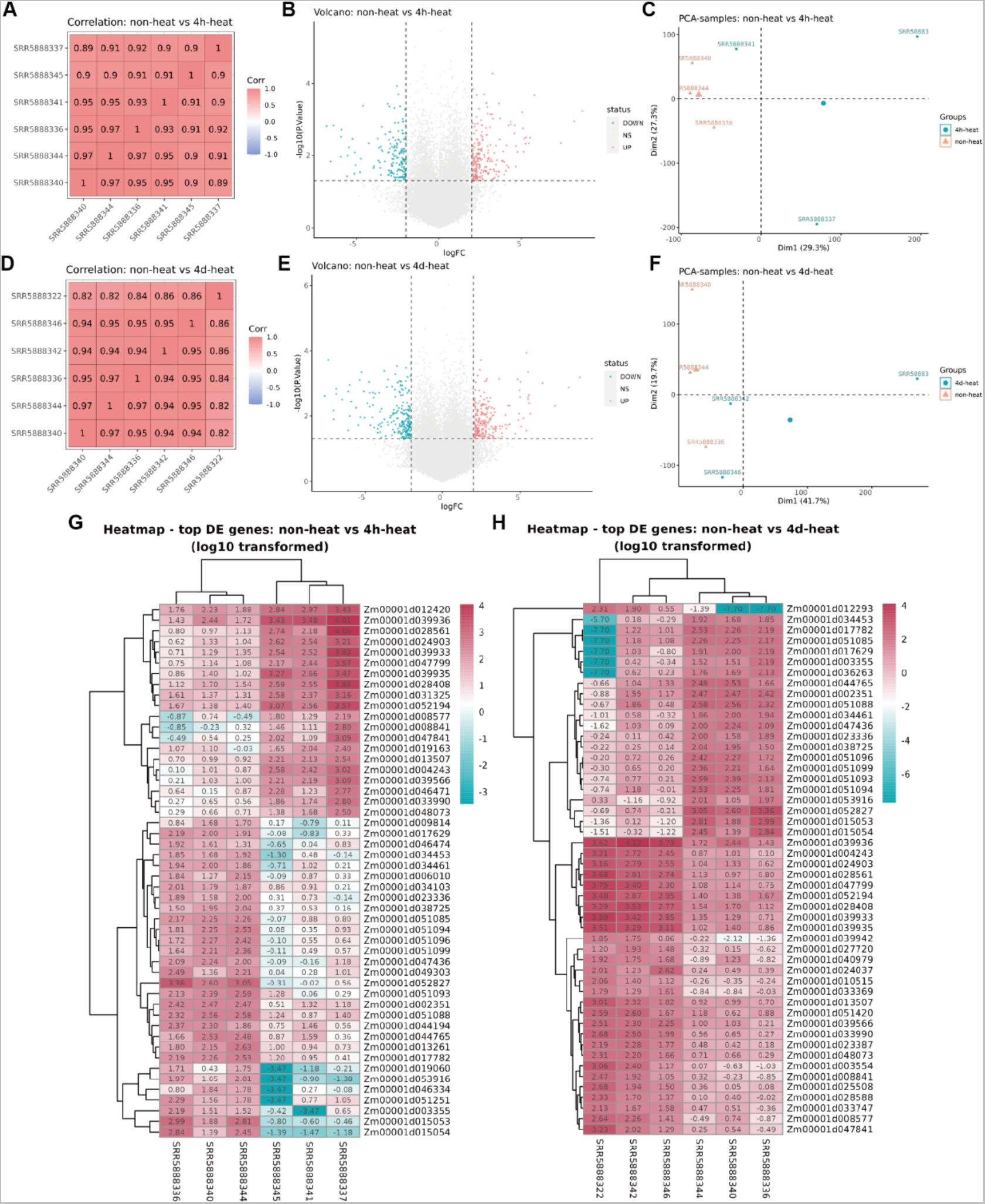
Differentially gene expression (DGE) analysis case study regarding heat stress conditions in *Zea mays* (0h/4h and 0h/4d). **(A)** Gene expression level correlation plot for all samples in the 0h/4h comparison. **(B)** Volcano plot for the 0h/4h comparison. **(C)** Sample PCA plot for the 0h/4h comparison. **(D)** Gene expression level correlation plot for all samples in the 0h/4d comparison. **(E)** Volcano plot for the 0h/4d comparison. **(F)** Sample PCA plot for the 0h/4d comparison. **(G)** Heatmap of expression level (TPM) regarding the top 50 differentially expressed genes in the 0h/4h comparison. **(H)** Heatmap of expression level (TPM) regarding the top 50 differentially expressed genes in the 0h/4d comparison.

**Supplementary Figure S3.**
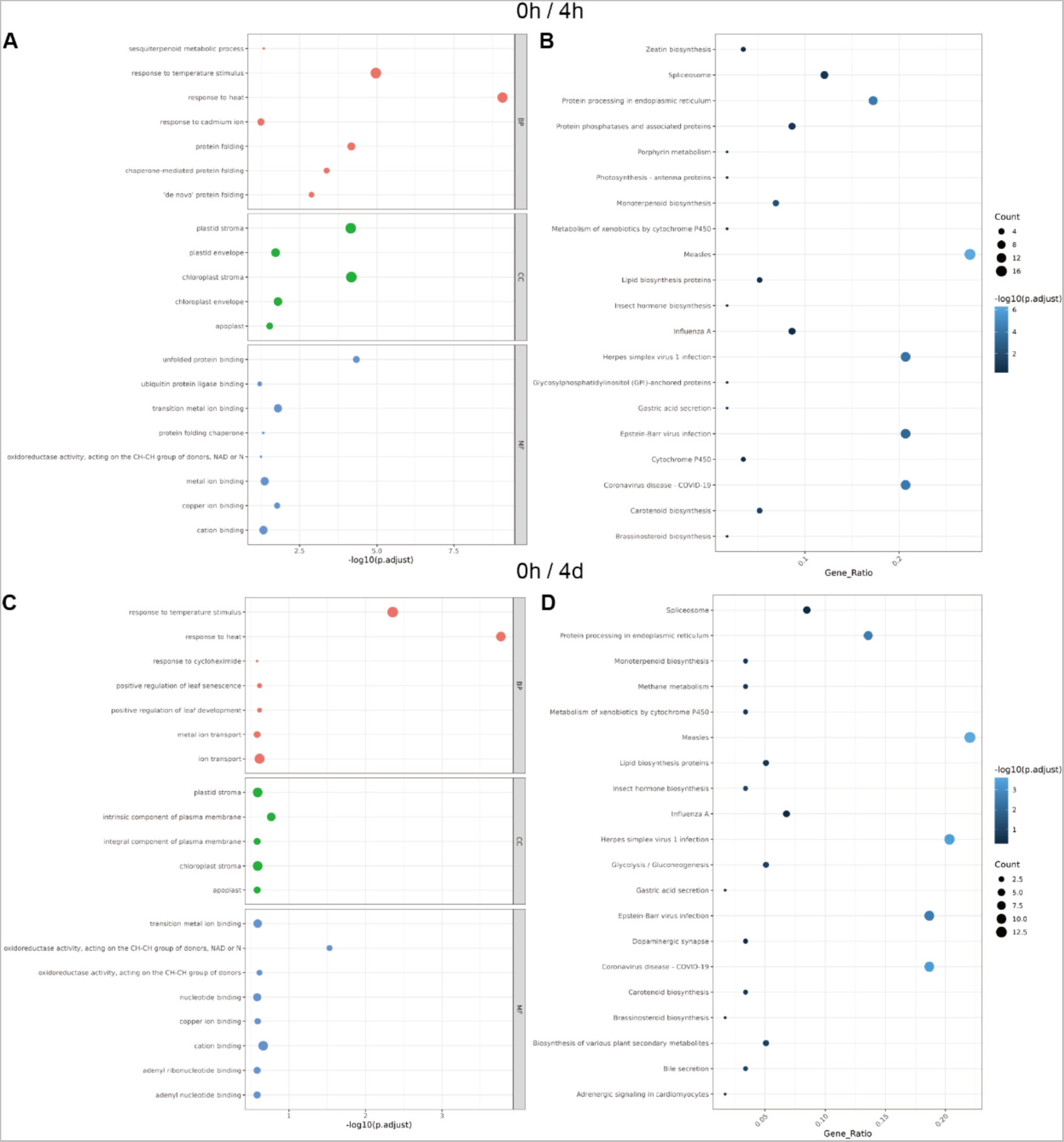
Functional enrichment of the differentially expressed genes (DEGs) regarding heat stress conditions in *Zea mays* (0h/4h and 0h/4d). **(A)** GO enrichment plot of differentially expressed genes in the 0h/4h comparison. **(B)** KEGG enrichment plot of differentially expressed genes in the 0h/4h comparison. **(C)** GO enrichment plot of differentially expressed genes in the 0h/4d comparison. **(D)** KEGG enrichment plot of differentially expressed genes in the 0h/4d comparison.

**Supplementary Figure S4.**
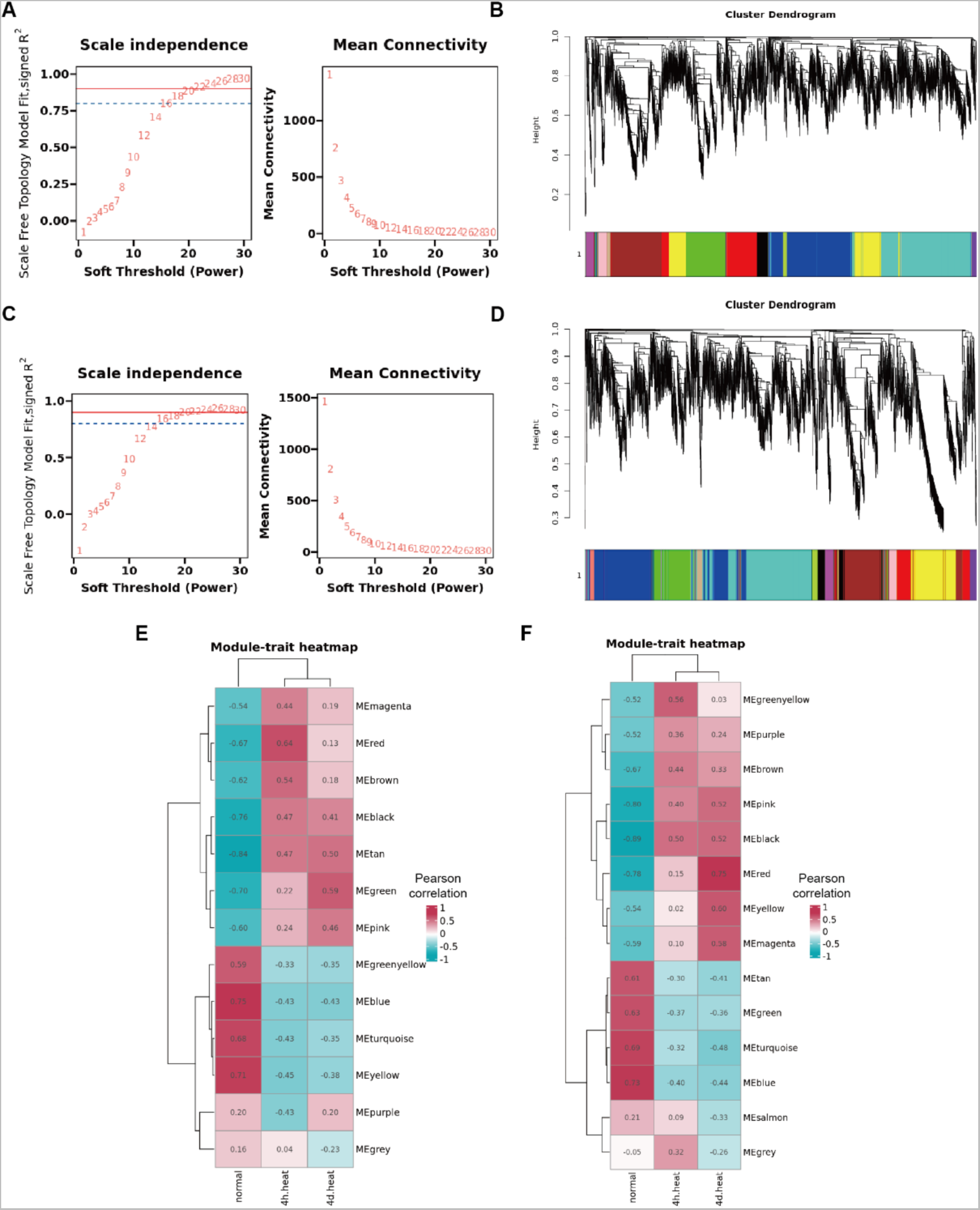
Weighted gene co-expression network construction and module-treatment correlation analysis of the top 4,000 differentially expressed genes (DEGs) regarding heat stress conditions in *Zea mays* (0h/4h and 0h/4d). **(A)** Scale independence and mean connectivity plot of the top 4,000 DEGs in the 0h/4h comparison. **(B)** Cluster dendrogram regarding the co-expression network constructed upon the top 4,000 DEGs in the 0h/4h comparison. **(C)** Scale independence and mean connectivity plot of the top 4,000 DEGs in the 0h/4d comparison. **(D)** Cluster dendrogram regarding the co-expression network constructed upon the top 4,000 DEGs in the 0h/4d comparison. **(E)** Module-trait correlation analysis heatmap between expression level of top 4,000 DEGs in the 0h/4h comparison and different heat stress treatments. **(F)** Module-trait correlation analysis heatmap between expression level of top 4,000 DEGs in the 0h/4d comparison and different heat stress treatments.

**Supplementary Figure S5.**
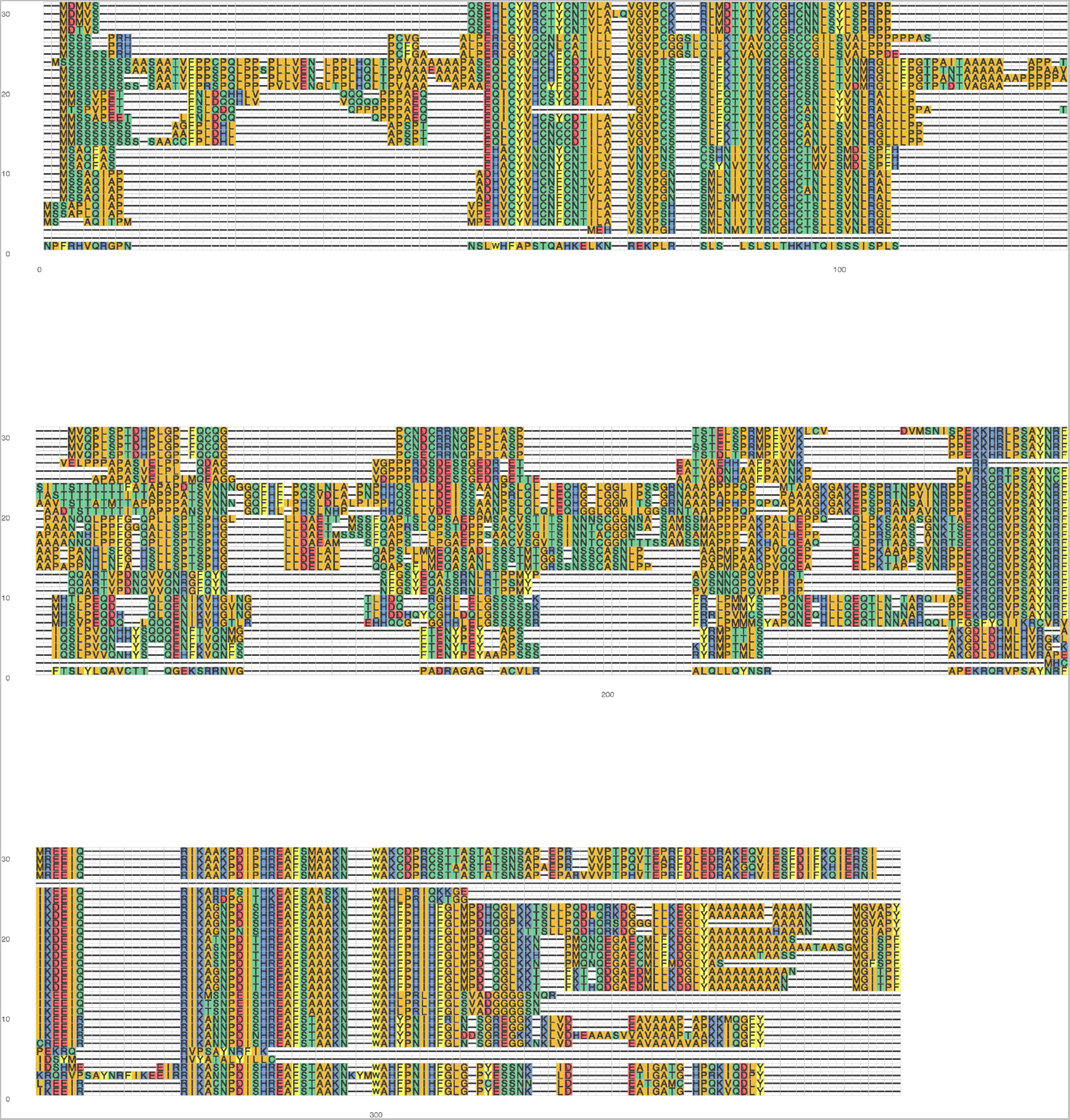
Protein multiple sequence alignment (MSA) plot of identified *YABBY* family candidates in *Zea mays*, *Sorghum bicolor* and *Coix lacryma-jobi*.

**Supplementary Figure S6.**
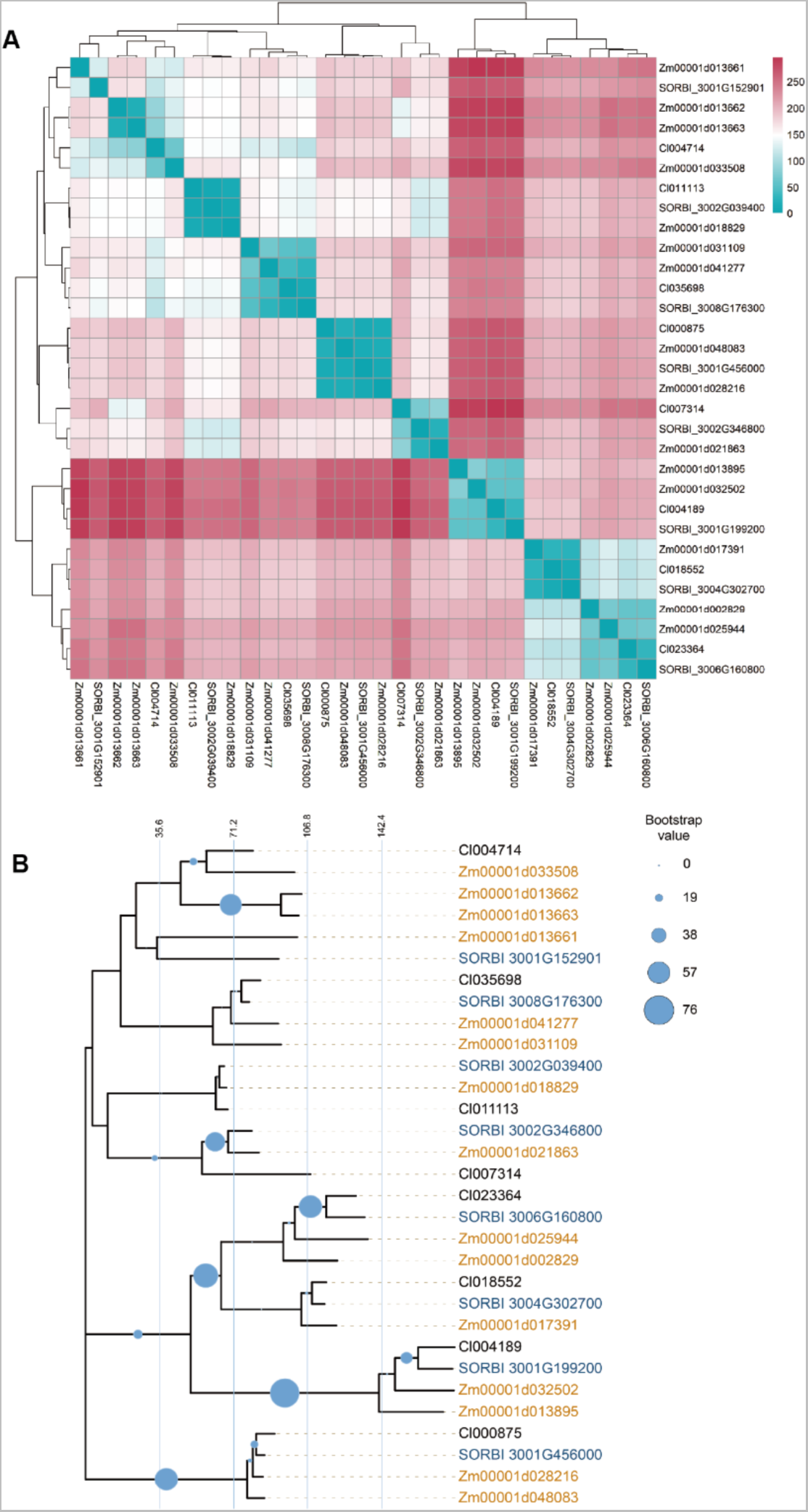
Pairwise protein sequence distance heatmap (Kimura 2-parameter method) and neighbor-joining phylogenetic tree of identified *YABBY* family candidates in *Zea mays*, *Sorghum bicolor* and *Coix lacryma-jobi*. **(A)** Pairwise protein sequence distance heatmap. **(B)** Neighbor-joining phylogenetic tree.

**Supplementary Figure S7.**
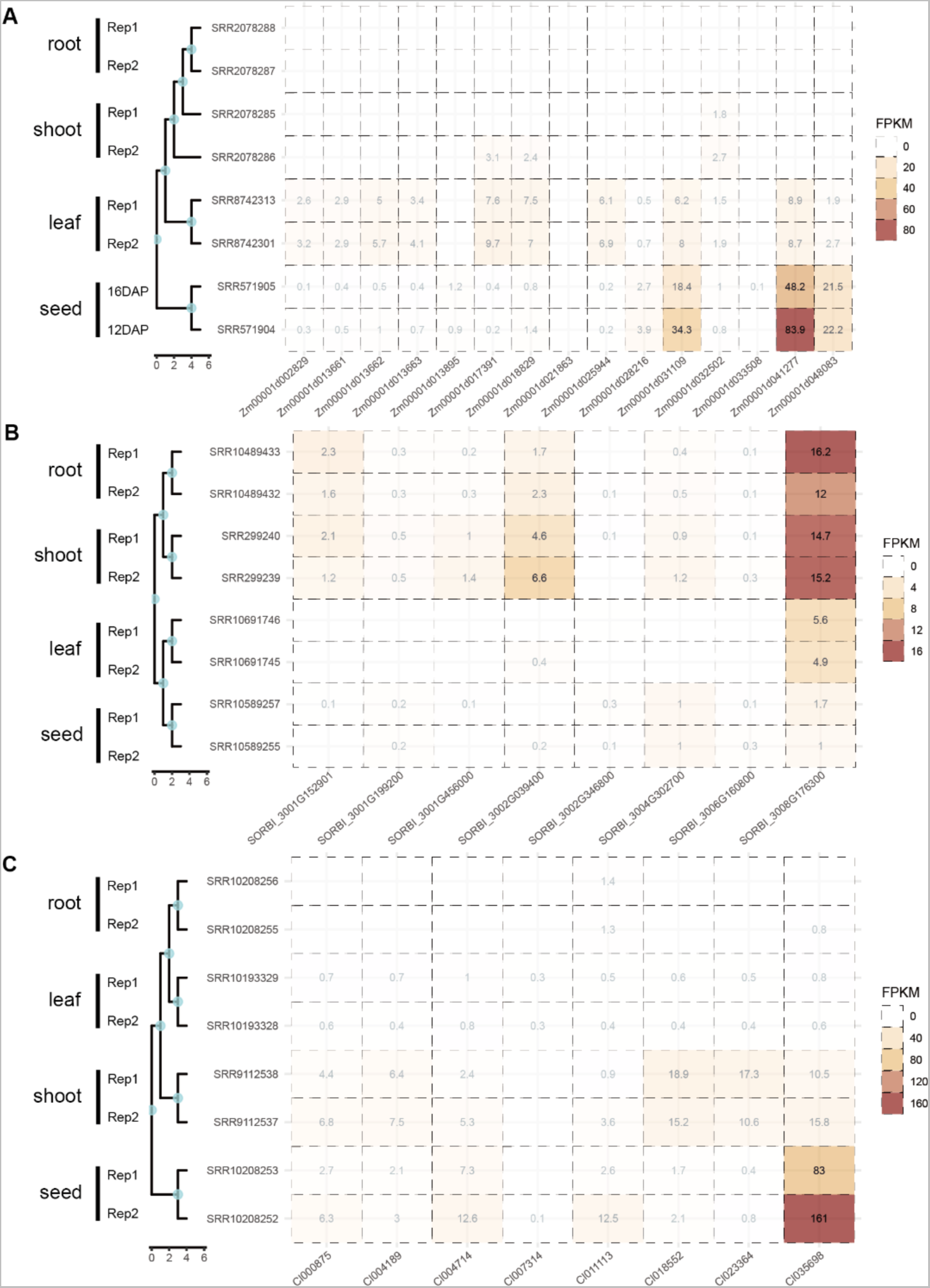
Expression heatmap of identified *YABBY* family candidates in *Zea mays*, *Sorghum bicolor*, *Coix lacryma-jobi* in root, shoot, leaf and seed tissues, each with 2 biological replicates (in FPKM). **(A)** Expression heatmap of identified *YABBY* family candidates in *Zea mays*. **(B)** Expression heatmap of identified *YABBY* family candidates in *Sorghum bicolor*. **(C)** Expression heatmap of identified *YABBY* family candidates in *Coix lacryma-jobi*.

